# Dual mechanism estrogen receptor inhibitors

**DOI:** 10.1101/2020.07.06.189977

**Authors:** Jian Min, Jerome C. Nwachukwu, Sathish Srinivasan, Erumbi S. Rangarajan, Charles C. Nettles, Valeria Sanabria Guillen, Yvonne Ziegler, Shunchao Yan, Kathryn E. Carlson, Yingwei Hou, Sung Hoon Kim, Scott Novick, Bruce D. Pascal, Rene Houtman, Patrick R. Griffin, Tina Izard, Benita S. Katzenellenbogen, John A. Katzenellenbogen, Kendall W. Nettles

**Affiliations:** State Key Laboratory of Biocatalysis and Enzyme Engineering, Hubei Collaborative Innovation Center for Green Transformation of Bio-Resources, Hubei Key Laboratory of Industrial Biotechnology, School of Life Sciences, Hubei University, Wuhan 430062, China; Department of Chemistry, Cancer Center and University of Illinois at Urbana-Champaign, Urbana IL 61801, USA; Department of Integrative Structural and Computational Biology, The Scripps Research Institute, Jupiter, FL 33458, USA; Department of Molecular and Integrative Physiology, Cancer Center and University of Illinois at Urbana-Champaign, Urbana IL 61801, USA; Department of Oncology, Shengjing Hospital of China Medical University, Shenyang, Liaoning Province, 110022, P. R. China; Department of Molecular Medicine, The Scripps Research Institute, Jupiter, FL 33458, USA; Omics Informatics LLC, 1050 Bishop St. #517, Honolulu, HI, 96813, USA; Precision Medicine Lab, 5349 AB Oss, The Netherlands

## Abstract

Efforts to improve estrogen receptor-a (ER)-targeted therapies in breast cancer have relied upon a single mechanism, with ligands having a single side chain on the ligand core that extends outward to determine antagonism of breast cancer growth. Here, we describe inhibitors with two ER-targeting moieties, one of which uses an alternate structural mechanism to generate full antagonism, freeing the side chain to independently determine other critical properties of the ligands. By combining two molecular targeting approaches into a single ER ligand, we have generated antiestrogens that function through new mechanisms and structural paradigms to achieve antagonism. These dual-mechanism ER inhibitors (DMERIs) cause alternate, non-canonical structural perturbations of the receptor ligand-binding domain (LBD) to drive antagonism of proliferation in ER-positive breast cancer cells and in allele-specific resistance models. Solution structural and coregulator peptide binding analyses with DMERIs highlight marked differences from current standard-of-care, single-mechanism antiestrogens. These findings uncover an enhanced flexibility of the ER LBD through which it can access non-consensus conformational modes in response to DMERI binding, broadly and effectively suppressing ER activity.

**Significance Statement:** To address the unmet clinical need for effectively suppressing estrogen receptor (ER) activity with both de novo resistance and in advanced ER-positive breast cancers that are resistant to standard-of-care antiestrogens, we have developed dual-mechanism ER inhibitors (DMERIs) that employ two distinct ER targeting moieties. These DMERI elicited non-canonical structural perturbations of the receptor ligand-binding domain and stabilized multiple antagonist sub-states within the dimer to generate highly efficacious antagonism of proliferation in ER-positive breast cancer cells and in allele-specific resistance models. This work reveals new conformational modes by which the activity of ER can be effectively suppressed to block breast cancer proliferation.

## Introduction

The estrogen receptor-α (ER) plays a critical role in breast cancer where it functions as a major driver of tumor growth in ~70% of breast cancers. The suppression of ER function with endocrine therapy is initially quite effective, either by inhibiting estrogen production with aromatase inhibitors (AIs) or by blocking ER activity with antiestrogens (1, 2). However, many ER-positive breast cancers recur in forms that have become resistant to standard-of-care AIs and/or antiestrogens, while de novo resistance also occurs. In these resistant cases, it is possible that antiestrogens of novel design might still prove effective because most of the tumor cells continue to express ER (3).

Currently, there are two types of approved antiestrogens. Tamoxifen (4) and its newer generation analogs, raloxifene, bazedoxifene and lasofoxifene, are called selective estrogen receptor modulators (SERMs) (5) due to their estrogenic activity in some tissues. SERMs all contain an additional aromatic ring that we call the E-ring (named with respect to the 4 rings of steroids that are lettered A-D, **Fig. S1A–B**). The E ring is attached to an aminoalkyl side chain via a two-carbon ether that exits from the ligand binding pocket and interacts on the receptor surface with a single H-bond at Asp351 (**Fig. S1A**); in this stabilized position, the side chain controls antagonism of breast cancer growth, as well as selective modulation of ER activity in other target tissues (6–10), all within a very tightly defined structural space on the receptor.

Fulvestrant, the only approved ER antagonist for treatment of tamoxifen or AI-resistant breast cancer (11), and other full antagonists such as RU 58,668, are termed selective estrogen receptor downregulators (SERDs), because they also reduce ER protein levels, although this effect may not be required for their antagonist activity (12–14). Both of these SERDs contain an extended terminal fluorine-substituted alkyl sulfinyl or sulfonyl sidechain (**Fig. S1B**), but suffer from poor pharmaceutical properties. Newer, orally active SERDs under clinical development have acrylate side chains (**Fig. S1C**), again having a 2-carbon linker but now to a carboxyl group (12, 15–20). Thus, both FDA-approved SERMs and potential oral SERD replacements for fulvestrant contain a single, carefully positioned aminoalkyl or acrylate side chain that through direct interaction with helix 12 (h12) in the ER ligand binding domain (LBD), moves it to disrupt the surface binding site for transcriptional coactivators that drive proliferative gene expression, thus operating by a *direct antagonism* mechanism of action.

Here, we take a different approach to block the activity of ER by combining within one ligand two distinct molecular elements that disrupt the conformation of the LBD: In addition to side chains typically used to effect *direct antagonism*, we add bulky chemical groups that cause *indirect antagonism* by distorting structural epitopes inside the receptor ligand binding pocket. We show here that indirect antagonism independently drives full antagonism, enabling the direct antagonist side chain to adopt new structural and functional roles that are associated with newly identified non-canonical conformations of the receptor. Coregulator peptide binding and structural studies highlight marked differences from current standard-of-care, single-mechanism antiestrogens. Thus, these dual-mechanism estrogen receptor inhibitors (DMERIs) represent a new class of antiestrogens that generate full antagonism of proliferation in wild type ER-positive breast cancer cells and in a number of allele-specific resistance models. Our findings uncover an enhanced flexibility of the ER LBD through which it can access non-consensus conformation modes in response to DMERI binding that very effectively suppress ER activity.

## Results

### Oxabicycloheptene Sulfonamide (OBHS-N) Core Ligands Display Inherent Indirect Antagonism and Offer Two Sites for Direct Acting Side Chain Attachment to Formulate Dual-Mechanism ER Inhibitors (DMERIs)

Indirect antagonists produce a range of activity profiles by interfering in new ways with the docking of h12 across h3 and h5 of the ER, which is required for formation of the surface binding site for transcriptional coactivator complexes, called Activation Function-2 (AF-2). Starting from a bulky oxabicyclic scaffold that contain aromatic rings corresponding to the A and E rings of other ER ligands (**Fig. 1A, Fig. S1A–C**), we appended a sulfonamide linker to prepare a 7-oxabicyclo[2.2.1]heptene sulfonamide (OBHS-N) core scaffold, as illustrated by the parental OBHS-N compound **13** (**Fig. 1A**). Despite lacking the canonical side chains required for the direct antagonism of standard of care antiestrogens, we previously showed that **13** was a full antagonist SERD, equivalent to fulvestrant in inhibiting proliferation of breast cancer cells with WT ER (21). The sizable −SO_2_-N(CH_2_CF_3_)(*p*-Cl-phenyl) group in **13** shifted the position of h11 by 2.4 Å; this blocked the interaction between the N-terminus of h3 and the C-terminus of h11 (**Fig. S1D**) and disrupted the agonist binding site for h12 against h11 (**Fig. S1E**), resulting in antagonist activity indirectly, that is, without direct interaction with h12. Compounds in this parental series, however, were not effective against the constitutively active mutants of ERa that drive treatment resistant disease (**Fig. S1E**) (21, 22).

**Figure 1.**
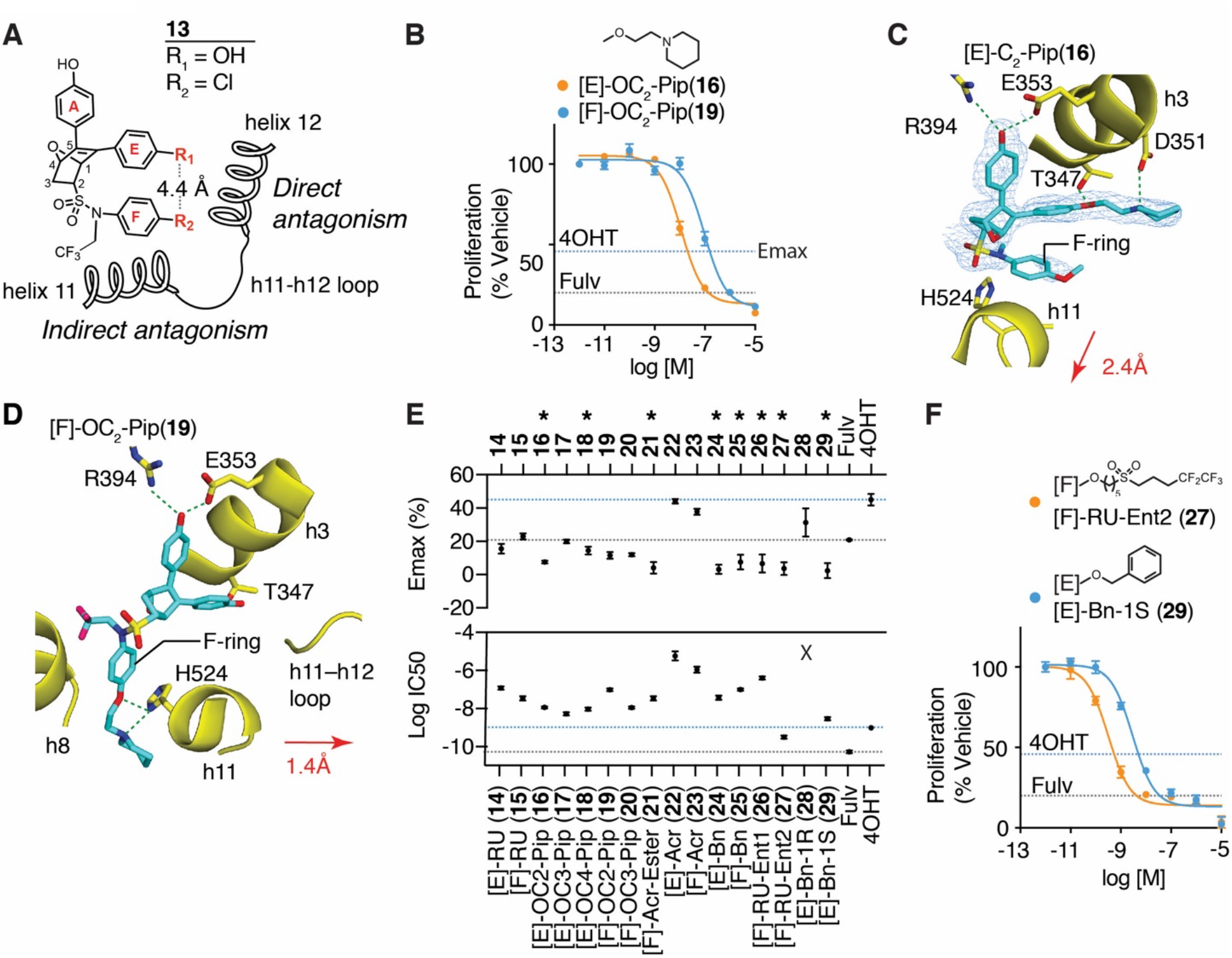
Dual-mechanism ER inhibitors fully suppress breast cancer cell proliferation. **A**) Chemical structure of the OBHS-N scaffold and the orientation of substituents R_1_ and R_2_, with respect to h11 and h12 in the ER-LBD (When R_1_ has a substituent, R_2_ is-OCH_3_ group; when R_2_ has a substituent, R_1_ is −OH for compounds **14-29**. See also **Fig. S2A-H**). **B**) Proliferation of MCF-7 cells treated for 5 days with 4OHT, fulvestrant (Fulv), or the indicated compounds. Datapoints are mean ± SEM, N= 6. M, Molarity. The horizontal lines indicate the maximum efficacy (Emax) for 4OHT and fulvestrant (Fulv). **C**) Structure of [E]-OC_2_-Pip (**16**)-bound ER LBD showed the E-ring substituted piperidine H-bonding to Asp351 in helix 3 (h3), while the F-ring shifts helix 11 (h11) by 2.4 Å compared to an agonist bound structure. 2F_o_-F_c_ electron density map contoured to 0.9 σ within 2 Å of the ligand. **D**) Structure of [F]-OC_2_-Pip (**19**)-bound ER LBD shows that its [F]-OC_2_-Pip side chain exits the ligand-binding pocket between h8 and h11, H-bonds to His524, and shifts h11 towards h12. **E**) Summary of dose response curves for compound inhibition of proliferation of MCF-7 cells, shown in **Figure S2A–H**. Datapoints are mean ± SEM, N= 6. Maximal efficacy, Emax. * indicates p_Adj_ < 0.05 (1-way ANOVA) for compounds with Emax > fulvestrant. **F)** Selected dose curves from **G)**. The lines indicate the maximum efficacy (Emax) for 4OHT and fulvestrant (Fulv)

The position of the E-ring inherent in the OBHS core closely matches the site from which the direct-acting antagonist side chains of standard of care SERMs and SERDs emanate. The sulfonamide substitutions that drive indirect antagonism of the OBHS-N compounds also contain another aromatic ring that we call the F-ring, and the flexibility of its linkage to the bicyclic ligand core enables the F-ring to extend in generally the same direction as the E-ring, in both cases towards h12 and the h11-h12 loop, typical targets of direct antagonism (**Fig. 1A**). Thus, the OBHS-N compounds provide two sites from which to launch a set of direct-acting antagonist side chains, the canonical E-ring site and the non-canonical F-ring site, the latter, which being 4.4 Å away, directs substituents into more unexplored structural space for ER antagonism (**Fig. 1A**). Our OBHS-N system thus allowed us to evaluate and compare two novel structure-based design approaches for ER antagonism: 1) combining indirect antagonism with the canonical direct antagonism emanating from substituents on the E-ring; and 2) combining direct and indirect antagonism from substituents emanating from the non-canonical F-ring location. To facilitate these comparisons, we chose in both cases to use the same set of substituents typically found in standard of care SERMs and SERDs, aminoalkoxy ethers, acrylates, and a Roussel-like extended alkyl sulfonyl group with a perfluorinated terminus. We also included some length variations in the aminoalkoxy ether groups, as well as ester precursors of the acrylate carboxylates and some benzyl ethers.

### Comparison of the Raloxifene Side Chain OC_2_-Piperidine Attached to the E-vs. F-Ring

We first compared OBHS-N compounds having the most traditional two-carbon aminoalkyl ether SERM side chain attached to the E-ring ([E]-OC_2_-piperidine (**16**)) or the F-ring ([F]-OC_2_-piperidine (**19**)) (**Supplemental Methods**). Both of these compounds inhibited the proliferation of MCF-7 breast cancer cells with greater efficacy (i. e., greater extent of proliferation suppression) than 4OHT (the active 4-hydroxy metabolite of tamoxifen) and equivalent efficacy to fulvestrant (**Fig. 1B**), but **16** was more potent than **19** (with 11 nM and 97 nM IC_50s_, respectively). We obtained X-ray crystal structures of these compounds in complexes with the ER LBD in the antagonist conformer, where h12 was displaced from the agonist position and flipped onto the activation function 2 (AF-2) surface to block coactivator binding (**Fig. S1F**).

As expected, the [E]-OC_2_-Pip side chain of **16** directly displaced h12 and formed a tight H-bond with h3 Asp351, whereas the unsubstituted F-ring in this ligand shifted h11 2.4 Å away from the position required for optimal agonist activity (**Fig. 1C, Fig. S1F**). To our surprise, the side chain of [F]-OC_2_-Pip (**19**) did not point towards h12, but instead exited between h8 and h11, where it is stabilized by H-bonds with h11 His524 (**Fig. 1D, Fig. S1G**). In this orientation, the F-ring attached antagonist side chain in **19** pushes h11 towards the agonist position of h12 by 1.4 Å, from which it is also expected to prevent formation of the agonist conformer. This novel, non-canonical orientation of an ER side chain also results in antagonism, which, though less potent than that of [E]-OC_2_-Pip (**16**), still fully inhibited breast cancer cell proliferation (**Fig. 1B**). Thus, the ER LBD complex with the F-ring-substituted OBHS-N compound **19** represents a new form of indirect antagonism, which was possible due to the presence of two rotatable bonds in the sulfonamide that allowed the F-ring to adopt multiple positions relative to the core (**Fig. S1H**). Examples of this were observed again in our series of crystal structures described below, and likely contributes to the unusual activity profiles of these ligands.

### Dual-Mechanism Inhibitors Produce Full Antagonism of Breast Cancer Cell Proliferation Irrespective of Their Side Chain Structure and Site of Substitution

To find other novel structural conformations that might arise from combining direct and indirect antagonism, we explored a wider range of side chains on both the E- and F-rings (**Fig. S2A–H**), including the one found in RU 58,668 ([E]-RU (**14**); [F]-RU (**15**)) and the acrylate found in orally active SERDs ([E]-Acr (**22**); [F]-Acr (**23**)). We also explored new side chains, including piperidines with longer 3- or 4-carbon linkers ([E]-OC_3_-Pip (**17**)), ([E]-OC_4_-Pip (**18**)), and ([F]-OC_3_-Pip (**20**)), as well as an acrylate-ester ([F]-AcrEster (**21**)) and simple benzyl substitution (([E]-Bn (**24**)) or ([F]-Bn (**25**), these latter ones being available from synthetic intermediates.

In terms of MCF-7 antiproliferative activity, aside from a few low potency compounds (acrylates **22** and **23** and the disfavored E-ring benzyl **28**), all of the E- and F-ring substituted compounds profiled as full antagonists (**Fig. 1E, Fig. S2A–H**). We consider this to be representative of the broad structural tolerance that results in full antagonism for ligands of the DMERI class through an expanded array of non-canonical conformations of their ER complexes, described below. Notably, 8 of the compounds showed maximal efficacy (Emax) significantly greater than fulvestrant (**Fig. 1E**, asterisks at top).

While the most of the side chains induced full antagonist, they differed in potency by almost four logs of IC_50_ values, demonstrating that with DMERI, the side chain can be used to optimize potency separately from effects on Emax. Because all of the OBHS-N compounds are racemates, we used chiral HPLC to resolve two of the compounds **15** and **24**, and we found that one enantiomer of each (**27** from **15**, and **29** from **24**) accounted for essentially all of their affinity and cellular activity. These preferred enantiomers (**27** and **29**) had very good antiproliferative IC_50s_ of 0.3 and 3 nM, respectively (**Fig. 1E, Fig. S2G–H**), and they were the focus of subsequent studies.

To verify on-target mechanism of action, we inhibited cell growth with a subset of the compounds and showed full pharmacological reversal with increasing doses of estradiol (**Fig. S2I**). We also showed that the compounds completely antagonized E2-induced expression of the ERa-target gene, *GREB1* (**Fig. S2J**), and notably, the compounds also had no effect on proliferation of MDA-MB-231 cells, a triple-negative breast cancer cell line that lacks ER (**Fig. S2K**), again supporting ER specificity.

### Atypical Side Chains Perturb the ER LBD Helix 12 Conformation

To further understand ligand-dependent effects on receptor structure, we compared X-ray crystal structures of ERa LBD complexes with dual-mechanism inhibitors and other antagonists. In the LBD, the [F]-OC_3_-Pip (**20)** longer side chain exited towards h12 and in doing so also shifted h11 by 1.6 Å to induce indirect antagonism (**Fig. 2A**). Here, the piperidine head group made van der Waals contacts with Trp383 in the same location where Pro535 in the h11-h12 loop typically resides in contact with Trp383 (**Fig. 2B**). This **20**-bound ER structure differed from those stabilized by SERMs such as raloxifene, as h12 was shifted 2.6 Å towards the C-terminus of h11, allowing Leu539 to directly contact the piperidine group of **20** (**Fig. 2B–C**). Unlike traditional SERM side chains that are stabilized by H-bonding, or the rigid acrylates of SERDs, the longer side chain of **20** is flexible with many degrees of freedom, suggesting that the shift in h12 is driven by both the shift in h11, which pulls on the h11-h12 loop, and additionally by F-ring the position of the atypical side chain.

**Figure 2.**
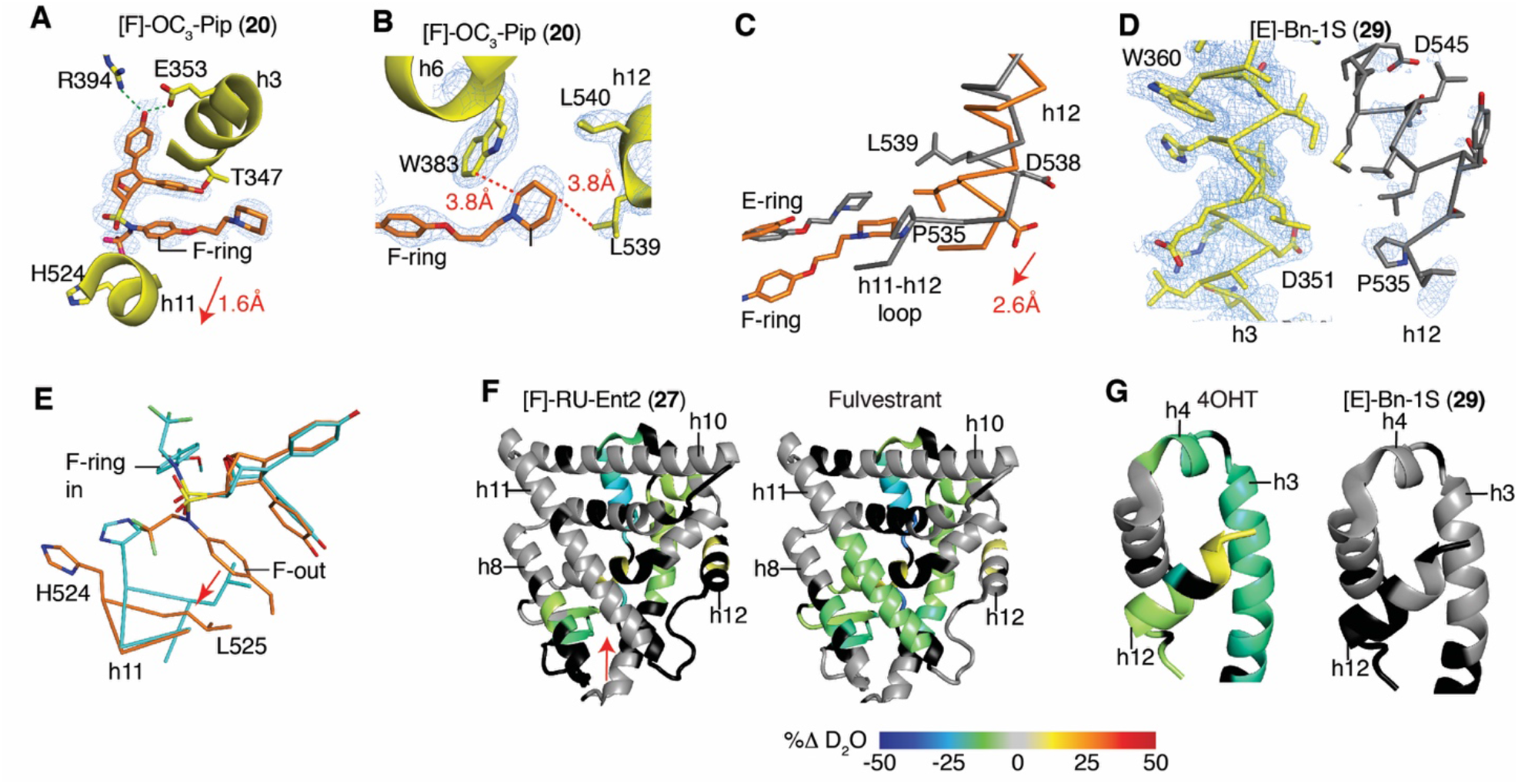
Dual mechanism inhibitors destabilize helix 12 of ERα. **A**) Structure of the ER LBD bound to [F]-OC_3_-Pip (**20**). 2F_o_-F_c_ electron density map contoured at 1.0 σ shows the **20** F-ring facing outward between h3 and 11 towards h12, shifting h11 1.6 Å compared to an agonist bound structure. The 2F_o_-F_c_ electron density map is contoured at 0.9 σ within 2 Å of the ligand. **B)** Structure of ER with **20** shows that **20** is stabilized by contacts with Trp383, which stabilizes the altered conformer of h12 by contacting L539. **C**) The structures of the ER LBD bound to **20** (coral) or raloxifene (gray) were superimposed, showing the 2.6Å shift of h12 to contact the piperidine ring of **20**. **D**) Structure of the ER LBD with [E]-Bn-1S (**29**) shows that h12 could not be modeled in 2 of 4 subunits due to poor electron density. The A chain of h3 (yellow) is shown with the B chain superimposed (gray) to show the expected location of h12, which was not modeled. The 2F_o_-F_c_ electron density map is contoured at 1.0 σ. **E**) Structure of [F]-AcrEster (**21**)-bound ER showing different ligand binding positions in the dimeric subunits. The A and B chains were superimposed and colored blue or coral. **F–G**) Changes in ER-Y537S H/D exchange compared to the apo receptor. ER-Y537S LBD was incubated with the indicated ligands and then assayed for exchange of amide hydrogens with deuterium over time, as measured by mass spectrometry. See also **Fig. S4**

The structure of the [E]-Bn (**29**)-bound LBD showed even more dramatic effects on h12, with very weak electron density where h12 was expected to be positioned, demonstrating that h12 was disordered (**Fig. 2D**). This is important, as the original two-position model of h12 (7) (**Fig. S1D** vs **F**) does not account for how certain antagonists recruit transcriptional corepressors. Structural and biochemical data indicate that the disordering or displacement of h12 renders a more open or accessible AF-2 surface, which is required for binding of a longer 3 helical peptide motif found in corepressors (8–10, 23) to support a more complete antagonism of proliferation.

In our structural studies, we noted that a number of the DMERI LBD complexes were ligand-induced conformational heterodimers (**Fig. 2B**), where genetically identical monomers bind the same ligand, but adopt different conformations in the context of the dimer (**Fig. S3**). This suggests that the ligand-induced shift in h11 of the dimer interface alters the conformer of the other monomer to favorably bind the second ligand differently. Many of the structures reveal the F-ring facing outward in one monomer and facing inward in the other monomer, associated with a smaller shift in h11 (**Fig S3C-E**). The [F]-acrylate (**23**) and [F]-acrylate ester (**21)** side chains differentially H-bonded to the N-terminus of h3 (**Fig. S3A**-**B**), which likely explained their widely different potencies. These effects were ligand selective (**Fig S3E**), and thus not driven by crystal packing. In addition to enforcing non-canonical receptor conformations, these ligands thus also stabilize an ensemble of both direct and indirect antagonist conformers, often within the same dimer.

### Hydrogen Deuterium Mass Spectrometry Reveals Alternate ER Conformers in Solution

To validate the destabilizing effects of the ligands on the ER LBD in solution, we examined the dynamics of secondary structural elements through mass spectrometry analysis of the exchange of amide hydrogens for deuterium using mass spectrometry (HDX-MS) (24). These included **30,** the high affinity enantiomer of the parental compound **13 (Fig. 1A**), the F-ring-substituted Roussel side chain compound **27**, and the E-ring substituted benzyl compound **29**. While 4OHT and fulvestrant stabilized the C-terminal half of h11 proximal to the ligands, the parental OBHS-N **30** and the dual-mechanism inhibitors **27** and **29** did not, consistent with indirect antagonism directed at h11 (**Fig. 2F, Fig. S4A**). All of the compounds stabilized h3 and h4 in the AF-2 surface, except for **29**, the compound that destabilized h12 in the crystal structure (**Fig. 2G**, **Fig. S4B**). With **27**, the extended hydrophobic RU side chain may directly contact the AF-2 surface to stabilize its secondary structural elements, as was seen with the fulvestrant analog ICI 164,384 in ERβ, which is the only available crystal structure of members of this class of SERDs (25) (**Fig. S4C**). With the other compounds, the stabilization of the AF-2 surface is likely through h12 binding to the AF-2 surface in the inactive conformer (**Fig. S1F**) (6, 7), highlighting the ability of **29** to destabilize h12 in solution and in the crystal structure. These studies demonstrate that with indirect antagonism, the shifts in h11 destabilize or reposition h12 of the ER LBD, allowing the side chains to have distinct roles in stabilizing alternate conformers of h12.

### DMERI Show Ligand Selective Activity Profiles in Other Cellular Contexts

It was striking that the DMERI as a class showed similarly robust Emax for inhibition of growth of wild type ERa-driven breast cancer (**Fig. 1B, E, F, Fig. S2A-H**) but showed a diversity of structural effects on the receptor (**Fig. 2, Fig. S3**). To probe whether the side chains supported different effects on SERM- or SERD-like activities of the ligands, we tested several compounds for effects on degradation of ERa, and found that while the parental indirect antagonists are SERDS (21), direct antagonist side chains determined whether compounds displayed SERM- or SERD-like properties (**Fig. 3A**). [E]-Bn (**24**) and the higher affinity enantiomer of [F]-RU, [F]-RU-Ent2 (**27**) were efficient ER degraders (**Fig. 3A**). These effects were reversed by 4-hydroxytamoxifen (4OHT) or the proteasome inhibitor MG132, demonstrating on-target mechanism of action through proteasomal degradation (**Fig. S5A**). In contrast, the compounds with piperidine side chains were more SERM-like, with either minimal effects on receptor stability (**14**, **17**, and **19**) or showing some stabilization of the receptor (**16** and **20**, **Fig. 3A**). [E]-Bn (**24**) required different ER domains for ligand-dependent degradation than seen with Fulvestrant (**Fig. S5B**), demonstrating that they have different effects on receptor structure.

**Figure 3.**
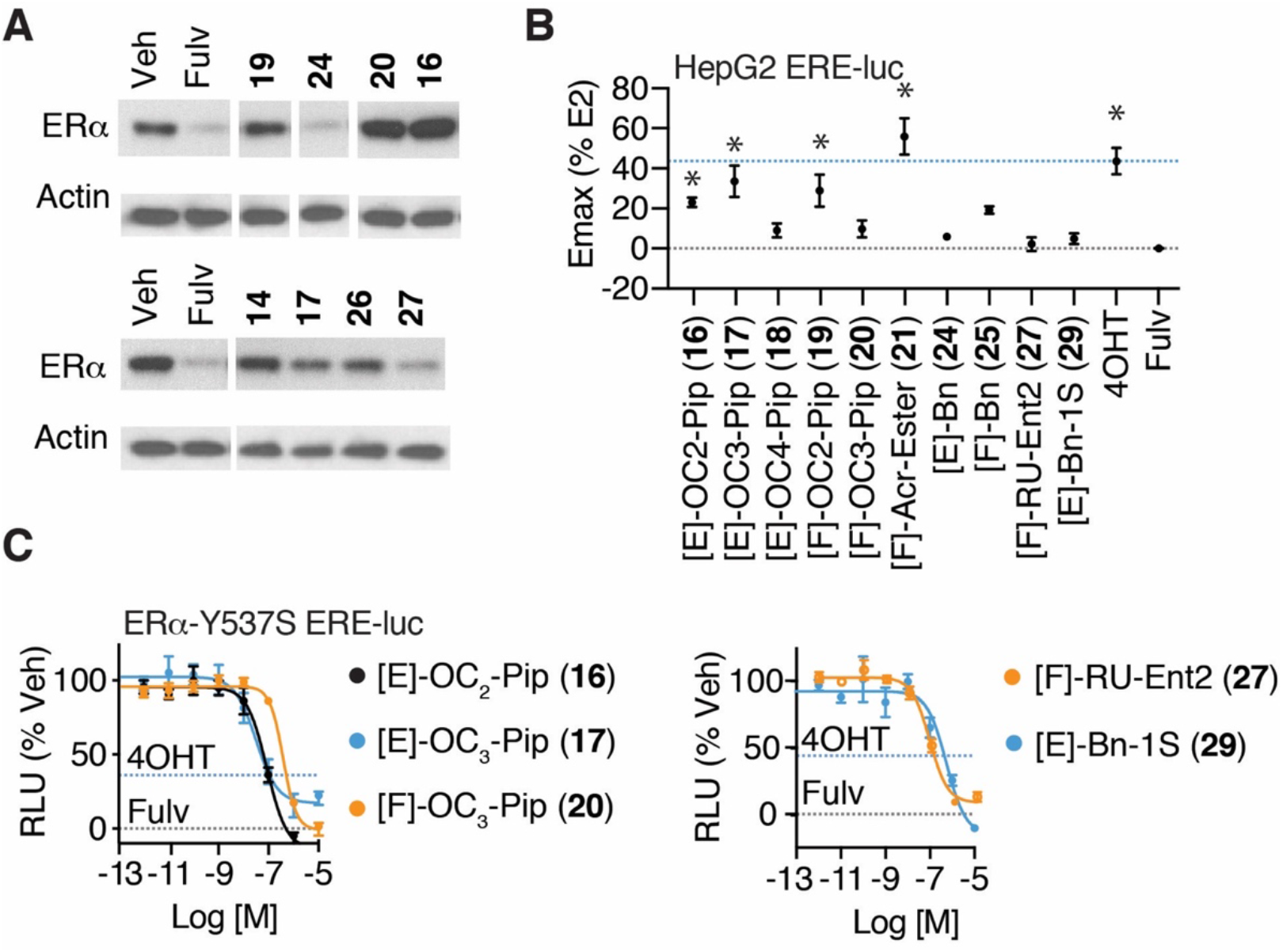
SERM and SERD properties of DMERI ligands. **A**) ER and β-actin levels in MCF-7 cells treated with the indicated compounds for 24 h. Whole cell lysates were analyzed by Western blot. See also **Fig. S5AB** **B**) Summary of dose response curves. Datapoints are mean ± SEM, N= 3. *Significantly different from fulvestrant by 1-way ANOVA, Sidak’s test adjusted p-value (p_adj_) < 0.05. N = 3–6. Dose curves are shown in **Fig. S5C**. **C**) Dose response curves with the indicated ligands in 293T cells. Datapoints are mean ± SEM, N= 3.

To further probe for the cell-type selective activity, we tested the compounds in HepG2 liver cells, because estrogenic effects on liver metabolism are an important contribution to protection from metabolic diseases (26). In these cells tamoxifen displays significant SERM agonist activity through the amino-terminal AF-1 domain of ER (27). We found that some of the compounds with piperidine-containing SERM side chains (**16, 17, 20**) showed cell-type specific agonist activity, as did [F]-acrylate ester (**21**), while the degraders **24** and **27** were full antagonists (**Fig. 3B, Fig. S5C**). Thus, the dual-mechanism inhibitor approach can produce compounds with SERM- or SERD-like properties that are highly efficacious and contain non-canonical side chains.

To test whether the compounds were effective against the constitutively active Y537S-ER that drives treatment-resistant metastatic disease and reduces the potency of antiestrogens, we tested the compounds with this receptor. Several of the piperidines were very efficacious (**16-19**), as were the [F]-AcrEster and [F]-RU compounds (**21** and **27**) (**Fig. 1G–H, Fig. S6**), demonstrating that both SERM and SERD DMERIs overcome the limitations of the parental indirect antagonists in this resistance model, and that differences between the DMERI are apparent in other contexts than inhibiting the WT ER in breast cancer cells.

### Dual-Mechanism Inhibitors Induce Unique ER Solution Structures by Peptide Interaction Profiling

To probe more deeply for molecular insights into distinctive bases for the non-canonical activity of DMERIs, we examined the interaction of full-length ER-WT or ER-Y537S complexes of DMERIs versus reference agonist and SERM and SERD compounds with a library of 154 peptides using the Microarray Assay for Realtime Coregulator-Nuclear receptor Interaction (MARCoNI) as a probe for solution structure (12, 28). Hierarchical clustering of the FRET data describing interaction of ER bound to 19 compounds x 154 peptide interaction profiles is shown in **Fig. 4A**, demonstrating a clustering of E2-induced peptide interactions (*Cluster 3*) that were strongly dismissed by 4OHT, fulvestrant, and the full antagonist SERDs, GDC-0810 and AZD9466 (*Cluster 1 vs 3*). We found individual peptides that showed some specificity, including PRDM2 (amino acids 948–970), which was recruited by 4OHT, NCOA1 (amino acids 737–759), which was selectively dismissed by the three SERDs compared to 4OHT, and NRIP1 (amino acids 805–831), which was not dismissed by fulvestrant (**Fig. 4B**). While it is possible to identify individual peptides that are selective for ER bound to these compounds (16, 29), most of the peptide interactions showed identical responses to the ligands in *Cluster 1*, with Pearson correlations (*r*) ≥ 0.90 between ligand-dependent peptide interaction profiles (**Fig. 4B**), highlighting the structural similarities of single-mechanism inhibitors including SERMs and SERDs.

**Figure 4.**
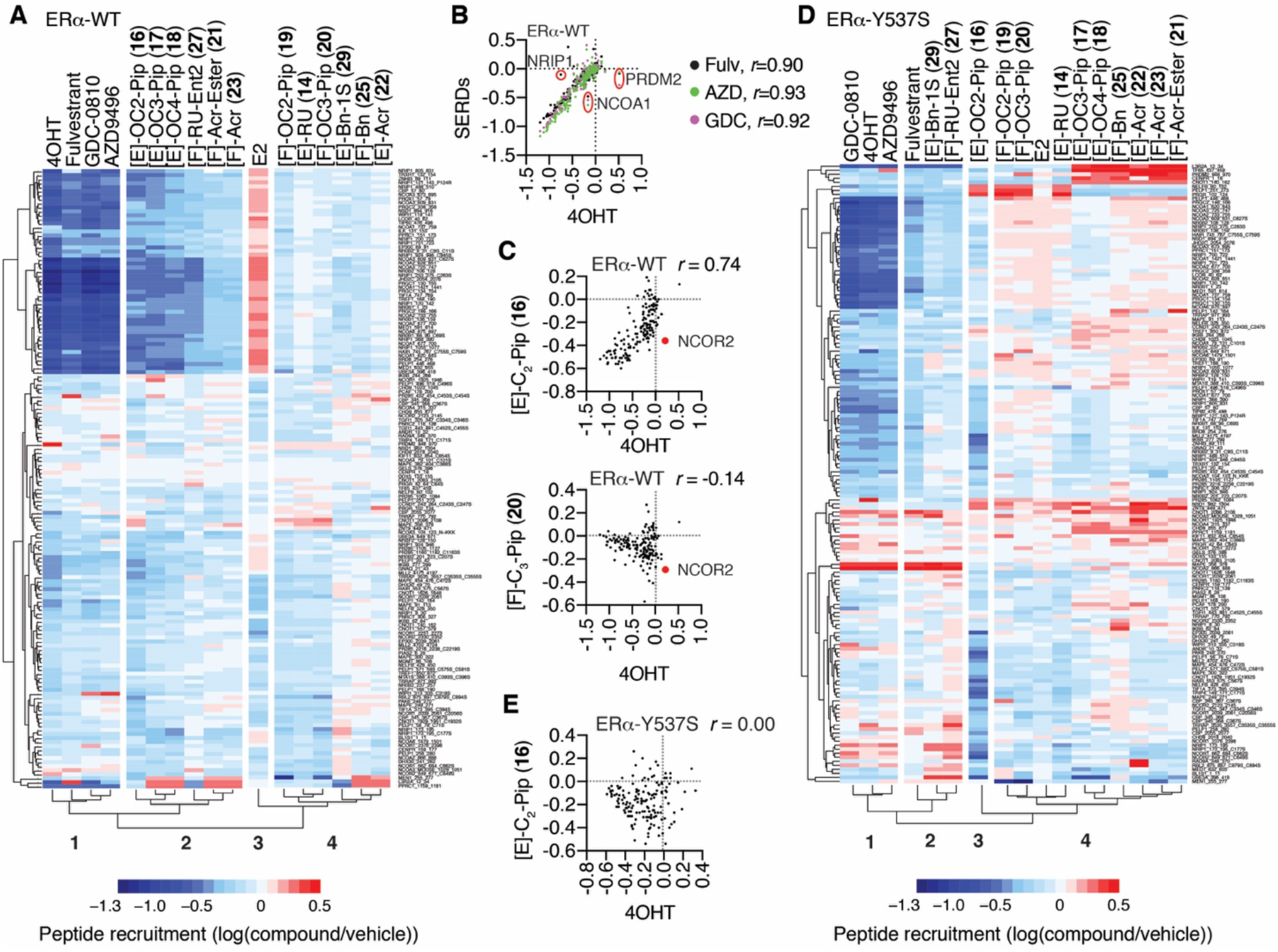
Dual-mechanism inhibitors promote conformations of ER that are distinct from traditional single mechanism inhibitors. **A**) Hierarchical clustering of MARCoNI (Realtime Coregulator-Nuclear receptor Interaction) FRET assay for interaction of full-length, wild type (WT) ER with 154 peptides derived from nuclear receptor-interacting proteins and the indicated ligands. **B-C**) MARCoNI Pearson correlations for 4OHT vs the indicated ligands. Fulvestrant (Fulv), GDC-0810 (GDC) or AZD9496 (ADZ). *r* = Pearson correlation ligand vs 4OHT **D**) Hierarchical clustering of MARCONI data with ER-Y537S and the indicated ligands. **E**) MARCoNI Pearson correlations for 4OHT vs [E]-C2-Pip (**16**) with the ER-Y537S. *r* = Pearson correlation. See also **Fig. S7**

*Cluster 2* was in the same clade as *Cluster 1* and contained the E-ring piperidine-substituted compounds, as well as the compounds with F-ring Roussel and acrylate side chains, all of which showed dismissal of the E2-induced peptide interactions (**Fig. 4A**). However, *Cluster 2* also has many more unique peptide interactions than *Cluster 1*, reflected in lower Pearson correlations (*r* = 0.65–0.74 vs 4OHT). These included NCOR2 peptide (amino acids 649-671), which is derived from a protein with context-selective coactivator or corepressor activity (30, 31), and was dismissed by **16** (and **20** in *Cluster 4)* but not 4OHT (**Fig. 4C**). *Cluster 4* displayed peptide interaction patterns most different from the traditional antagonists, and included the compounds with F-ring piperidines, E-ring RU or acrylate side chains, and both the E- and F-ring substituted benzyl compounds. For example, **20** showed very little overlap in peptide interaction patterns with 4OHT and showed many peptides that were selective for **20** compared to 4OHT (**Fig. 4C, bottom**). Thus, the dual-mechanism inhibitors displayed a variety of different solution structural features that differentiate them from the traditional antagonists, all of which displayed highly similar interaction profiles.

The ER-Y537S mutation changed the clustering pattern of ligand-dependent peptide interactions. 4OHT, AZD9496, and GDC-0810 still clustered together and dismissed many of the same E2-induced peptide interactions (*Cluster 1*, **Fig. 4D**), while fulvestrant now clustered with [F]-RU-Ent2 (**27**) and [E]-Bn-1S (**29**) in *Cluster 2* from the same clade. Despite this clustering pattern, fulvestrant still displayed a higher Pearson correlation with 4OHT (**Fig. S7A**), as they strongly dismissed many of the peptides (**Fig. S7B–C**). The other major clade includes *Cluster 3*, which contains only [E]-OC_2_-piperidine (**16**), and *Cluster 4*, which contained all the remaining piperidine-containing compounds, the acrylates, as well as the [E]-RU (**14**) and [F]-Bn (**25**) compounds. The dramatic shift in peptide interaction patterns is underscored by **16**, which with the ER-Y537S displayed no overlapping effect on peptide interaction patterns with 4OHT (**Fig. 4E vs 4C**). These observations highlight the similarities in solution structures of ER-WT or ER-Y537S bound to the traditional direct antagonists but point to a range of novel solution structures for many of the dual-mechanism inhibitors, some of which are unique to the mutant ER-Y537S. Given that the surface structure of ER controls the recruitment of transcriptional coactivators and corepressors as conveyers of receptor activity, these unique conformations likely contribute to the distinct activity profiles of the dual-mechanism inhibitors.

### Activity of Dual-Mechanism Inhibitors in Allele-Driven Models of Antiestrogen Resistance

Approximately one-third of patients with recurrent ER+ breast cancers present with constitutively active ER mutations, including Y537S and D358G (28, 32–34), while de novo EGFR overexpression drives a worse outcome and tamoxifen resistance in a significant subset of newly presenting breast cancer patients (35, 36). To explore these two modes of endocrine therapy resistance, we overexpressed EGFR in MCF-7 cells (**Fig. S8**), which we compared to parental MCF-7 cells as well as those engineered to express ER-Y537S or ER-D538G (33). With these models we observed the expected loss of both potency and efficacy in response to 4OHT or fulvestrant, with the EGFR model showing a complete loss of response to 4OHT (**Fig. 5A**). Our two high potency enantiomers, **27** and **29**, and two ligands with SERM properties (**16** and **20**) that had shown unusual peptide binding (**Fig. 4**) and structural features (**Fig. 1C vs Fig. S3C, Fig. 2A–C**) were highly efficacious in blocking Y537S activity (**Fig. 3C**). All of the ligands showed reduced efficacy, but **27** showed better potency in the mutant ER models, while **20** and **29** showed slightly better potency in the EGFR resistance model (**Fig. 5B**). Despite their diverse side chains for direct antagonism, all of these compounds suppressed proliferation across resistance models, highlighting the important role of the dual mechanism for antagonizing ER actions.

**Figure 5.**
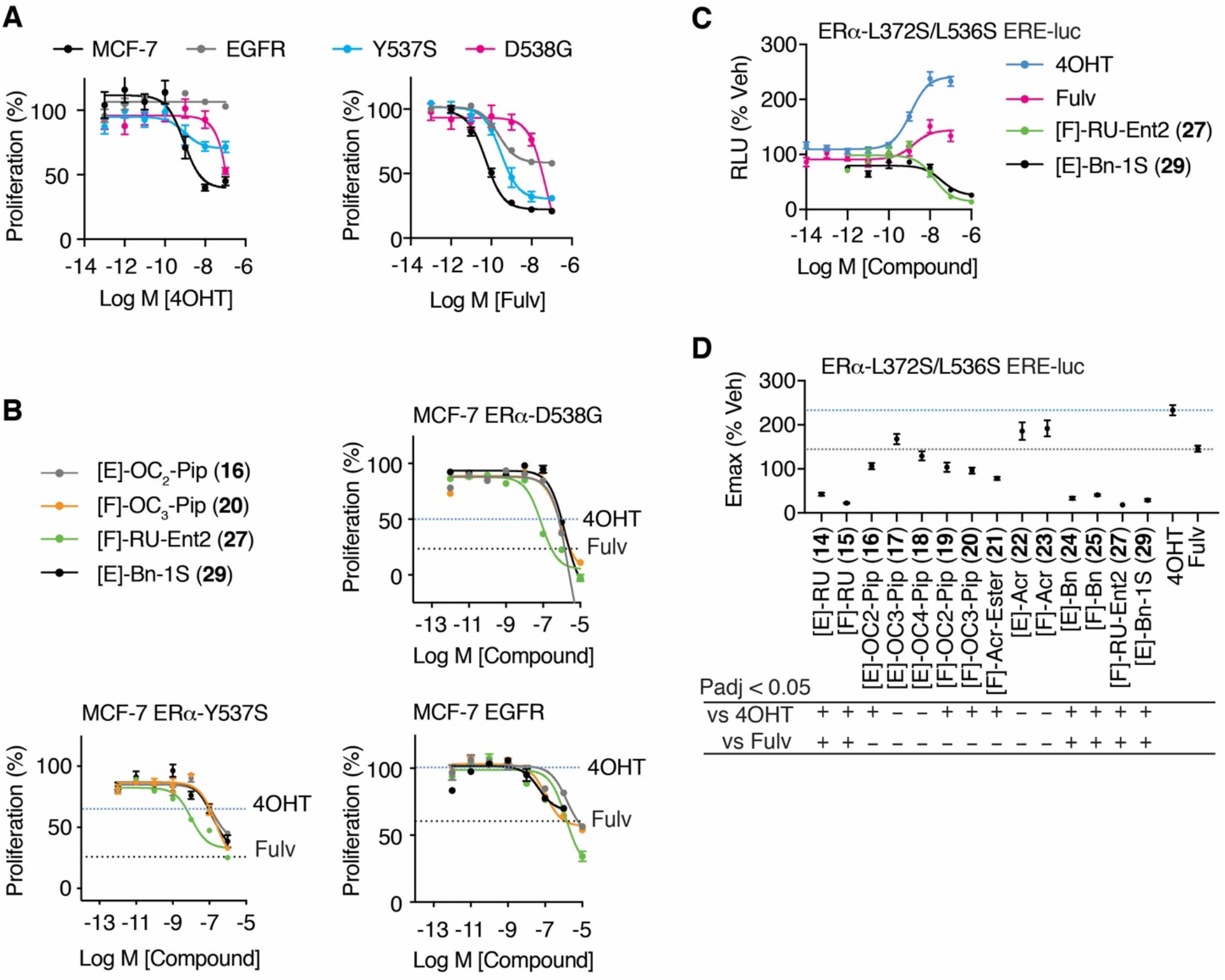
Activity of ligands in allele-specific models of tamoxifen resistance. **(A)** WT MCF-7 cells and **(A–B)** MCF-7 cells engineered to express the mutant ERα-Y537S or ERα-D538G, or overexpress EGFR, were treated with the indicated ligands for 5 days and analyzed for inhibition of cell proliferation. Fulvestrant (Fulv). N = 3. Dashed lines indicate Emax values for 4OHT or fulvestrant. See also **Fig. S8.** **C–D)** Structure-based model of tamoxifen and fulvestrant resistance. HepG2 liver cells were treated for 24 hr with the indicated ligands. N = 6, except for 4OHT and Fulvestrant where N=18. Data were analyzed by 1-way ANOVA. Data are mean ± S.E.M.

To further explore endocrine-resistance modes in breast cancer, we used a model we previously developed as the first structure-based design model of resistance for tamoxifen (27). Using the tamoxifen-bound ER structure to design L372S/L536S as a set of mutations to stabilize h12 as seen in that structure docked into the AF-2 surface, we found that these two mutations blocked binding of the NCOR1 corepressor to ER and enforced AF1-dependent SERM agonist activity for tamoxifen (21, 37). Mutations of L536 have since been identified in metastatic patient samples, highlighting its important role in regulating h12 dynamics (34, 38). Here, we demonstrate that this model also renders fulvestrant an agonist, in essence inverting the activity profile for these two standard-of-care antiestrogens from antagonists to agonists. In using this model for compound profiling (**Fig. 5C–D**), we found most of the compounds to be more efficacious than fulvestrant, including those with SERM like side chains, while the RU (**14**, **15**, **27**) and benzyl (**24**, **25**, **29**) side chain compounds were significantly more efficacious, almost completely blocking the AF1-driven activity that was enhanced by tamoxifen or fulvestrant (**Fig. 5C–D**). In this context, nearly all of the dual-mechanism ligands were significantly more efficacious than 4OHT (**Fig. 5D**). A number of the SERM like piperidine containing compounds had Emax values greater than fulvestrant where they did not stimulate activity (**16**, **19**, and **20**), while benzyl- and RU-substituted compounds were significantly better, strongly inhibiting the constitutive AF-1 driven activity (**Fig. 5C–D**).

## Discussion

In this work, we show dual-mechanism ER inhibitors (DMERIs) to function as a flexible chemical platform for the generation of ligands with tailored SERM or SERD like properties that are broadly efficacious across different breast cancer anti-estrogen resistance models, including a structure-based design model of tamoxifen and fulvestrant agonist activity. Probing of ER solution structures with a library of interacting peptides revealed that the DMERIs imposed unique solution structures that have important and favorable functional characteristics, an insight that was supported by HDX studies, whereas traditional single mechanism inhibitors—whether SERM or SERD—overall stabilized very similar structures. Our crystallographic structural analyses then demonstrated that these ligands induced unique perturbations to h11 and h12 to support their strong antagonism, including the formation of conformational heterodimers, where each monomer component of the receptor is in effect “reading” the same ligand in two different ways, an interaction that seems to be associated with the most efficacious DMERIs.

Our findings provide new insights beyond the traditional view of nuclear receptor allostery, which is based on single, direct-acting mechanism in which the ligand adopts a single pose to control the conformation of the protein (**Fig. 6A-C**) (39). The single-mechanism ligands select or induce lowest energy conformations of the receptor associated with specific activity profiles that can be active, inactive, or tissue selective (**Fig. 6A–D**). The targeting of multiple antagonist sub-states with DMERIs may provide a therapeutic targeting advantage similar to the effects of targeting multiple growth pathways with combination therapies, or the combined use of bazedoxifene and conjugated estrogens to achieve unique ER-mediated signaling characteristics (40).

**Figure 6.**
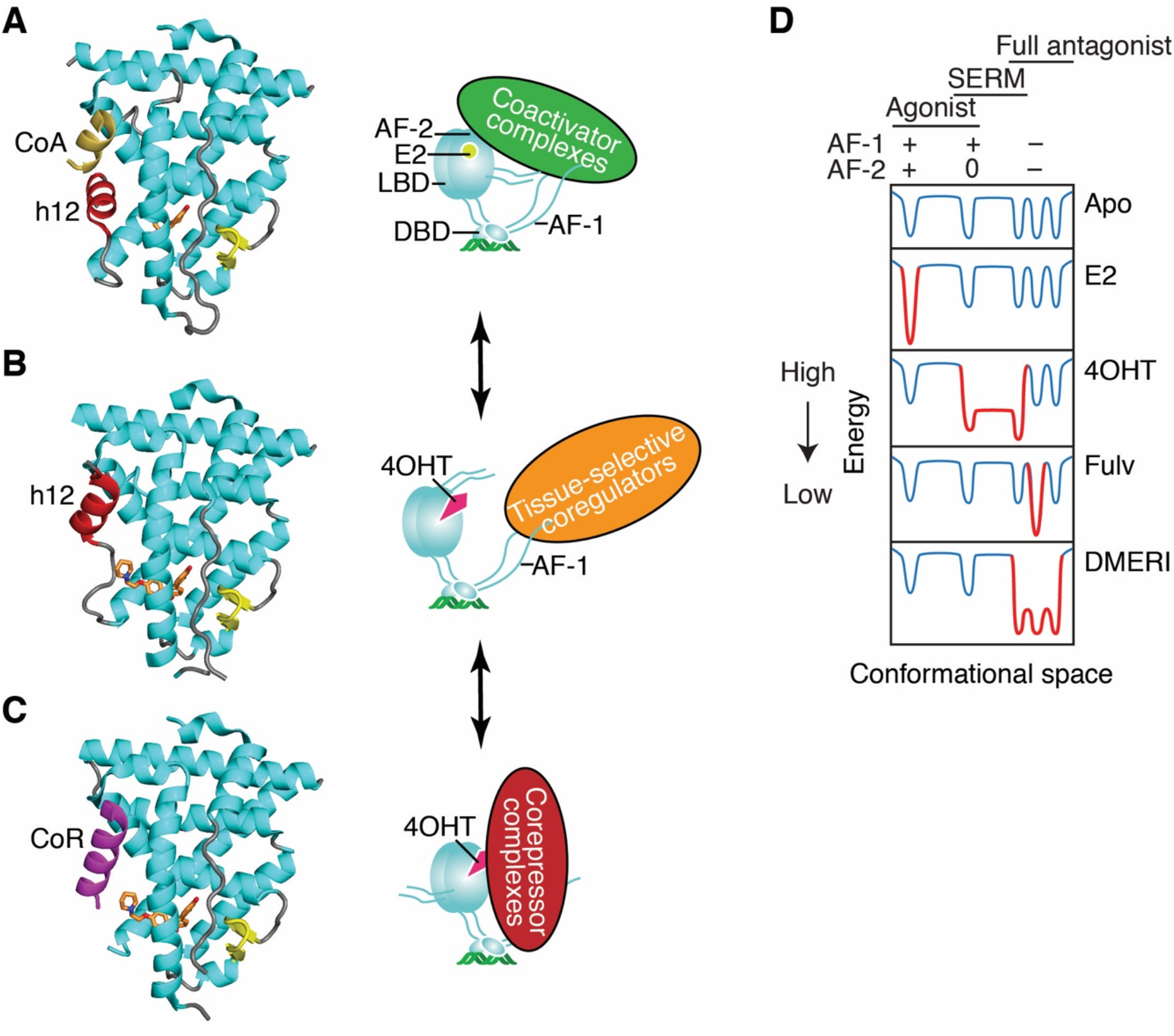
Ligand-dependent control of ERa-LBD conformation, and of ER coregulator recruitment and selection of activity states. **A**) The active LBD conformation (7). *Left*, ribbon diagram of the ER LBD bound to estradiol. Helix 12 (h12, colored red) forms one side of the coactivator binding site, shown here binding to a peptide from Steroid Receptor Coactivator-2 (CoA colored yellow) from pdb entry 3UUD. *Right*, Schematic of ERα bound to estradiol (E2), DNA, and a coactivator complex. With full agonists, the coactivator recruitment to the LBD surface, AF-2 (Activation Function-2) nucleates binding of multi-protein coactivator complexes to other domains including AF-1 (Activation Function-1). DBD, DNA binding domain. Steroid Receptor Coactivators (SRCs) 1–3 bind to both AF-1 and AF-2 through separate interactions. **B**) The transcriptionally repressive LBD conformation (9, 12, 25, 52). *Left*, ribbon diagram of the ER LBD bound to a corepressor peptide, colored violet. When h12 is disordered by an antagonist, the LBD can bind an extended peptide motif found in transcriptional corepressors(8) from pdb entry 2JFA. *Right*, Cartoon of ERa bound to 4-hydroxytamoxifen (4OHT) and a corepressor complex, repressing both AF-1 and AF-2 activity and mediating mediate chromatin compaction and inhibition of proliferative gene expression. **C**) The inactive LBD conformer (7, 27). *Left*, Ribbon diagram of the ER LBD bound to an antagonist. Antagonists can flip h12 (colored red) into the coactivator/corepressor binding site, rendering the LBD inactive by blocking both coactivator and corepressor binding to AF-2, from pdb entry 2QXS. *Right*, when h12 blocks both coactivators and corepressors from binding the LBD, the activity of AF-1 is cell-type specific. **D**) Energy diagram illustrating how ER ligands differ in stabilizing specific low energy receptor conformations associated with transcriptional activity (+), inactivity (0), or repression (–) that are being driven by the activity state of AF-2 or AF-1. The dips in the curves represent different LBD conformations associated with the three AF-1/AF-2 activity states shown at the top, *leftmost* being the active state (***a***), the *rightmost* representing substates of the repressive state (***b***), and the *middle* the inactive state (***c***). When a state is stabilized by a particular type of ligand, the curves become deeper, with gray changed to red; the barrier heights between states indicate the ease of dynamic interchange among the states or sub-states. The DMERI showed multiple mechanisms of antagonism, represented by the multiple favored repressor sub-states with reduced exchange barriers.

A key feature of the OBHS-N scaffold used here is that the indirect antagonism drives full suppression of ER activity, which we showed with the parental compounds lacking a side chain (21). This then enabled the added side chains to take on different functional roles. Since the development of tamoxifen in the 1970s and fulvestrant in the 1990s, the next generation SERMs and SERDs have directed either an aminoalkyl group to push on the h11-h12 loop or the acrylate unit to pull on it, leading to compounds with side chains that were localized around a very tight structural interface with ER. With DMERI, the h11-h12 loop was pulled indirectly via h11 to destabilize h12. This enabled a diversity of side chain activities from either the E-ring or F-ring in the DMERIs to dial back in SERM activity (**Fig. 3B**), bind directly to h12 to produce altered antagonist conformers (**Fig. 2B–C**), or produce efficacy even greater than fulvestrant by fully destabilizing h12 with full antagonists (**Fig. 1B**, **Fig. 2D, Fig. 2G**, **Fig. 5B–D**). This advance greatly expands the potential design principles for the ligand side chain that is not being used as the primary driver of antagonism and explains why we observed strong antagonism even when the side chain did not engage in the known modes of antagonism or was completely disordered. Thus, combining two chemical targeting approaches—direct and indirect antagonism—into a single ligand provides a flexible platform for new ER-directed therapies with different targeted signaling outcomes and broad efficacy across different treatment resistance models.

## Data and materials availability

All data needed to evaluate the conclusions in the paper are present in the paper and/or the Supplementary Materials. Compound profiling source data will be provided in a supplementary_data.xls upon acceptance. PDB coordinates will be released upon acceptance. Structures and map files are available to reviewers upon request from the Editor.

## Materials and Methods

### Cell Culture

MCF7, MCF7-ERα-Y537S, MCF7-ERα-D538G, HepG2, and MDA-MB231 cells were maintained in Dulbecco’s modified eagle medium (DMEM) supplemented with 10% fetal bovine serum (FBS). MCF7-ERα-Y537S, MCF7-ERα-D538G were a gift from Dr. Steffi Oesterreich, University of Pittsburgh Medical Center. The cells lines above were cultured with 1% penicillin/ streptomycin/ neomycin (PSN) antibiotics, 1% MEM non-essential amino acids, and 1% GlutaMAX (all from Gibco™ by Thermo Fisher Scientific), maintained at 37°C in a 5% CO_2_ incubator. Cells were tested regularly for mycoplasma contamination.

**Luciferase co-transfection assay and cell proliferation assay** are as previously described (21, 41) and detailed in **Supplemental Methods.**

### Quantitative RT-PCR (qPCR)

Total RNA was isolated using the RNeasy kit with on-column DNase I digest (QIAGEN). 4 μg total RNA samples were reverse-transcribed in 40 μl reactions using the High-Capacity RNA-to-cDNA™ Kit (Thermo Fisher Scientific, cat. no. 4387406). cDNA samples were analyzed by real-time PCR in triplicate 10 ul reactions using the 2X TaqMan^®^ gene expression master mix (Applied Biosystems™ by Thermo Fisher Scientific, cat. no. 4369016) with human *GREB1* (Hs00536409_m1) and *GAPDH* (Hs02758991_g1) expression assays. Relative mRNA levels were compared using the ΔΔCt method.

### Western Blot

Cells were lysed in ice-cold RIPA buffer (20 mM Tris pH 7.5, 150 mM NaCl, 1% NP40, 0.5% Sodium deoxycholate, 1 mM EDTA and 0.1% SDS). Protein samples were loaded on Any kD™ Mini-PROTEAN^®^ TGX™ Precast Protein Gels (Bio-rad, Hercules, CA) and transferred onto PVDF membranes (Thermo Scientific, Rockford, IL). The membranes were blocked with PBS-T + 5% nonfat dry milk and probed with primary antibodies overnight. The next day, the membranes were washed with with TBS-T, and incubated with HRP-conjugated probes (Santa Cruz Biotechnology) and developed using an ECL detection system (GE Healthcare Bio-Sciences, Pittsburg, PA).

### Antibodies and Probes

ERα (F-10) mouse mAb (1:1000, cat. no. sc-8002), ERα (H222) rat mAb (1:1000 dilution, cat. no. sc-53492), β-Actin (C4) mouse mAb (1:10,000 dilution, cat. no., sc-47778), HRP-conjugated mouse IgG kappa binding protein (cat. no., sc-516102), and HRP-conjugated goat anti-rat IgG antibody (cat no. sc-2006), were purchased from Santa Cruz Biotechnology, Inc.

### Macromolecular X-ray Crystallography

The ERα-L372S/L536S double-mutant ligand-binding domain (LBD, amino acid residues 298–554) was expressed in BL21 (DE3) E. coli cells, purified by IMAC using a Ni2+ column, dialysis, TEV digest, ion exchange, and size exclusion chromatography to remove the HA tag, as previously described(22). The purified LBD was co-crystallized with various ligands through sitting drop vapor diffusion method using trial gradients of 20% −25% (w/v) PEG 3350, 200 mM MgCl_2_, and pH 6.5–8.0, as previously described (37, 42). Data was collected at the Stanford Synchrotron Radiation Lightsource (Beamline: 12-2) and Advanced Photon Source (Beamlines: SER-CAT BM22, ID-22), both at temperature of 100 K and wavelength of 1.0 Å and scaled using AutoPROC (43) with the application of STARANISO (Globalphasing) to accommodate anisotropic diffraction. The structures were solved by molecular replacement of the starting model, PDB entry 2QXS, and then rebuilt and refined using the PHENIX software suite version 1.16 (44, 45). Ligand restraints were built on the PHENIX electronic Ligand Builder and Optimisation Workbench (46). Ligand docking was automated with LigandFit in Phenix and visually inspected using COOT version 0.8.9.2, as previously described (47, 48). New structures were further refined on the PDB-REDO server (49), before final refinement and validation in the PHENIX environment. Structures were analyzed using COOT and imaged using PyMOL (Schrodinger).

### MARCoNI coregulator interaction profiling

Microarray assay for real-time nuclear receptor coregulator interaction (MARCoNI) was performed as previously described (50). Hek293 cells were transfected with full length HA-tagged WT ERa or Y537S-ERa A PamChip peptide micro array with 154 unique coregulator-derived NR interaction motifs (#88101, PamGene International) was incubated with extracts from the 293T transfected cells in the presence of 10 μM compound or solvent only (2% DMSO, apo). Receptor binding to each peptide on the array was detected using fluorescently labeled HA-antibody, recorded by CCD and quantified. Per compound, three technical replicates (arrays) were analyzed to calculate the log-fold change (modulation index, MI) of each receptor-peptide interaction versus apo. Significance of this modulation was assessed by Student’s t-Test.

### Hydrogen-Deuterium Exchange (HDX) detected by mass spectrometry (MS)

Differential HDX-MS experiments were conducted as previously described with a few modifications(51), described in **Supplemental Methods**.

