## Supplemental Information for "Dual mechanism estrogen receptor inhibitors"

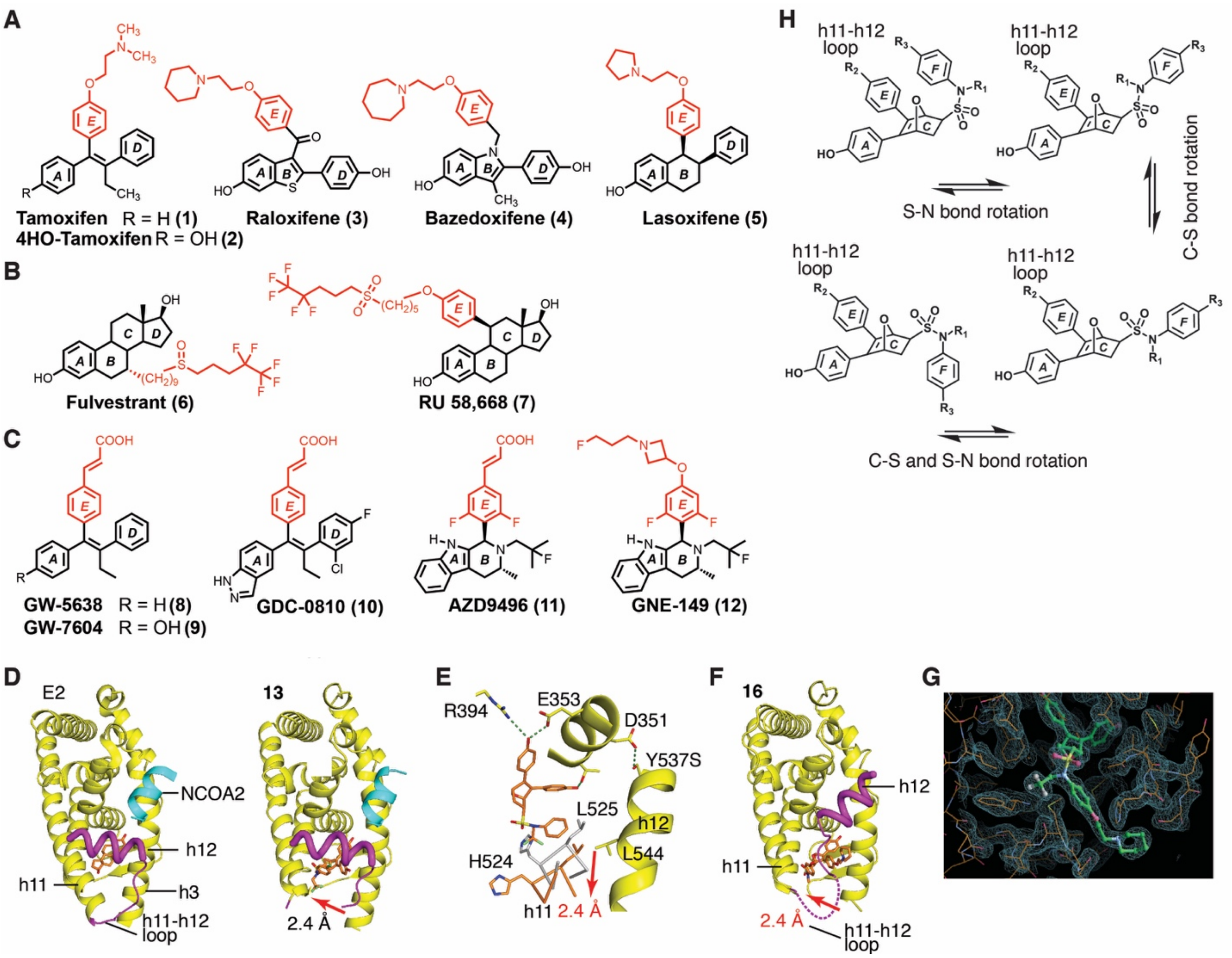


**Figure S1. Representative chemical structures and structural background**

**A**) Representative SERMs with basic side chains. Rings are lettered based on ERα binding relative to a steroid (A–D). Addition of another ring (the E-ring) to an agonist provides the traditional site for attaching antagonist side chains.

**B**) Full antagonists/SERDs with extended hydrophobic side chains.

**C**) Full antagonists/SERDs with an acrylic acid side chain.

**D**) Agonist and indirect antagonist/partial agonist structures. The ERα LBD is shown as ribbons with helix 12 colored purple and a peptide from the NCOA2 coactivator colored blue. When bound to the full agonist estradiol, (E2), the C-terminus of helix 11 (h11) and the N-terminus of h3 form direct contacts, providing a stable platform for the docking of h12 in the agonist conformer, enabling coactivator binding to the AF-2 surface. With the indirect antagonist OBHS-N **13**, h11 is shifted by 2.4Å away from h3, leading to the disruption of the end of h11 and the h11-12 loop.

**E**) Structure of **13** (yellow ribbons and coral sticks) superimposed on the structure of E2, bound to the Y537S-ERα LBD. The F-ring shift in h11 disrupts the favorable VDW packing of h12 L544 against L525. This destabilization of h12 is countered by the Y537S mutation which forms a tight h-bond with D351 in h3, which is why this ligand was ineffective in blocking activity of this mutant ER.

**F**) Structure of [E]-OC2-Pip (**16**) shows that the piperidine side chain flips h12 onto the AF-2 surface, while the F-ring shifts h11 by 2.4Å, leading to the h11-h12 loop being disordered.

**G**) 2Fo-Fc map contoured at 1σ shows the binding position of **16**.

**H)** Illustration of the conformational flexibility of the OBHS-N core ligands, which are capable of rotating around both the S-N and C-S bonds.

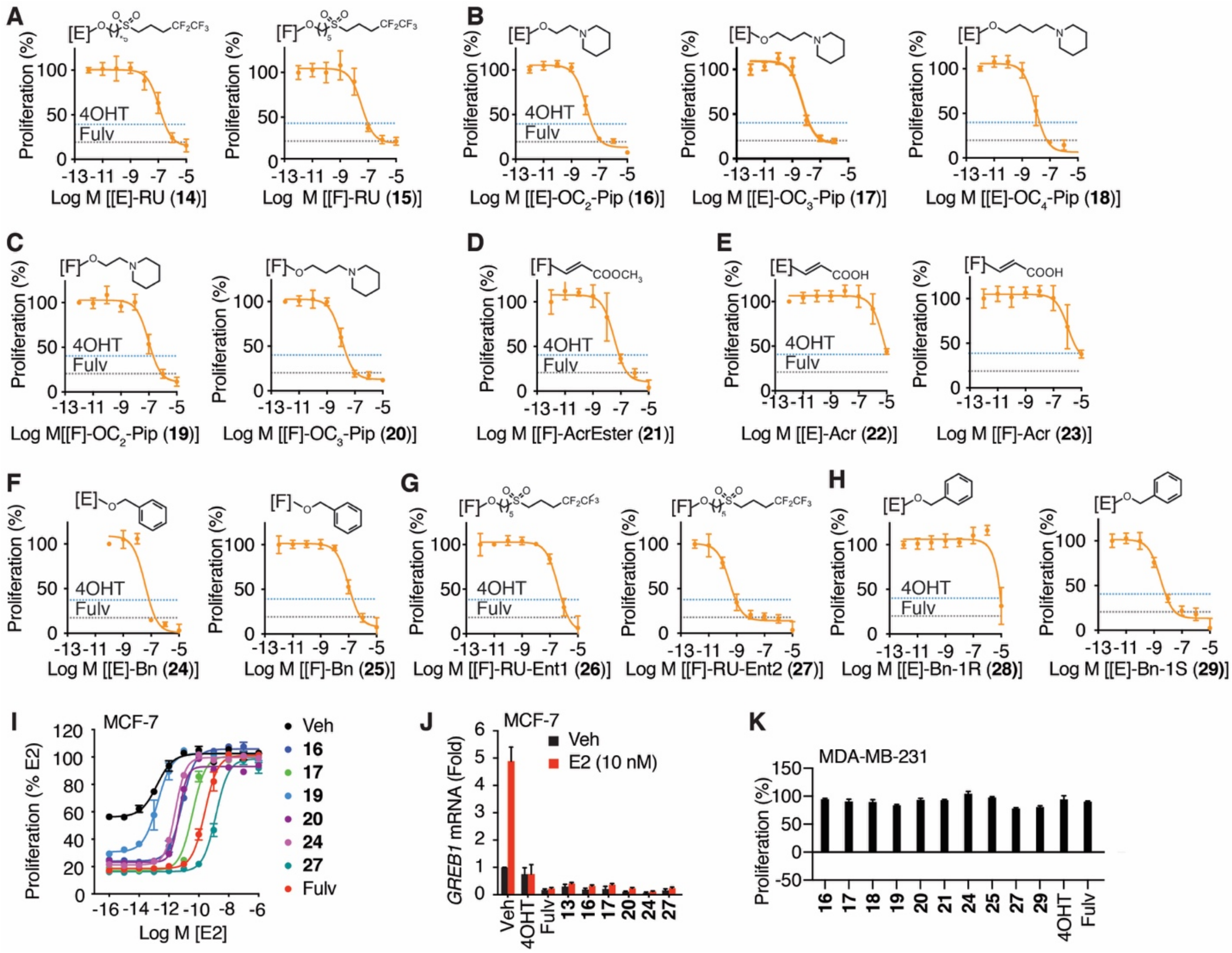

**Figure S2. Effects of dual mechanism inhibitors on breast cancer cell proliferation and their on-target mechanism of action.**

(**A–H**) MCF-7 cells were treated for 5 days with 4OHT, fulvestrant, or the indicated compounds. N= 6 x 8 doses.

**A**) Roussel (RU58,668) side chains attached to the E/F-ring.

**B–C**) Basic side chains.

**D**) Ester side chains.

**E**) Acrylic acid side chains.

**F**) Benzyl ether side chains.

**G**) Separate enantiomers of **15**.

**H**) Separate enantiomers of **24**. The absolute configuration was assigned based on the crystal structure of **29** bound to ERα.

**I**) Reversal of the antiproliferative effect of compounds by increasing concentrations of E2. MCF-7 cells were treated with the indicated compounds at 1 μM and cotreated with increasing doses of E2. Proliferation was measured after 5 days. N = 3.

**J**) Ligand-dependent suppression of E2-stimulated *GREB1* mRNA expression. MCF-7 cells were treated with 1 μM of the indicated compounds ± 10 nM E2 for 24 h and analyzed by qPCR. N = 3.

**K**) ERα negative MDA-MB-231 cells were treated with the indicated ligands at 1 □M for 5 days and analyzed for effects on cell number. N = 3

Data are mean ± SEM

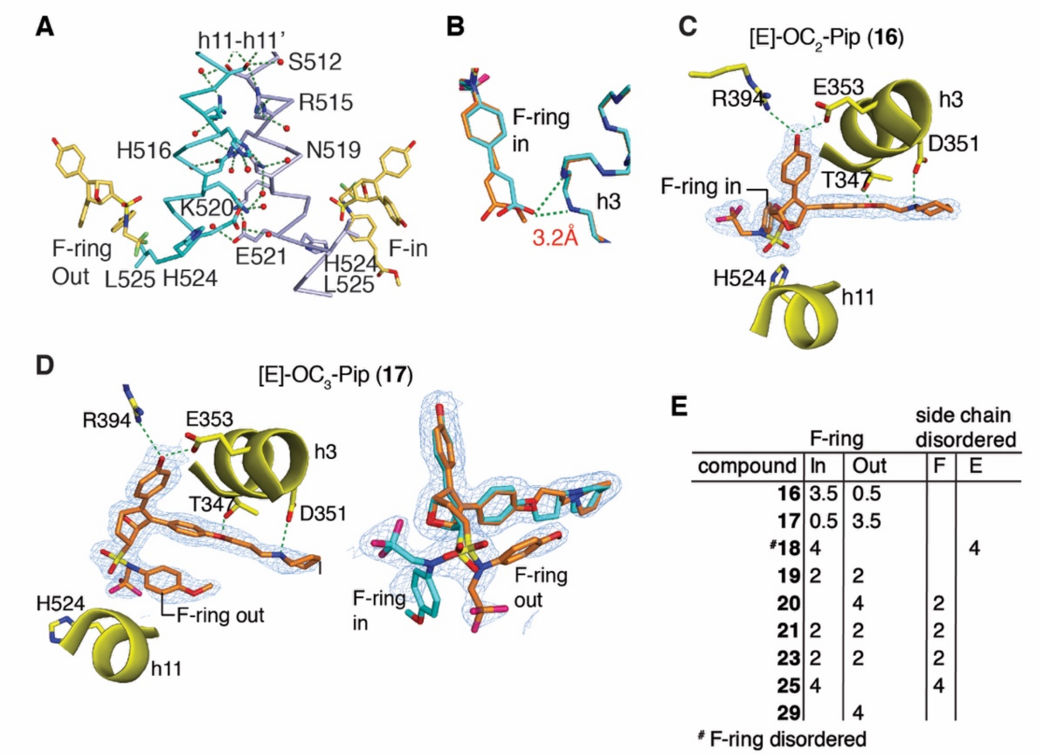


| | F-ring | | side chain disordered | |
|---|---|---|---|---|
| compound | In | Out | F | E |
| **16** | 3.5 | 0.5 | | |
| **17** | 0.5 | 3.5 | | |
| #**18** | 4 | | | 4 |
| **19** | 2 | 2 | | |
| **20** | | 4 | 2 | |
| **21** | 2 | 2 | 2 | |
| **23** | 2 | 2 | 2 | |
| **25** | 4 | | 4 | |
| **29** | | 4 | | |

### F-ring disordered

**Figure S3. Dynamic ligands that bind in more than one orientation drive multiple antagonist conformers and the formation of conformational heterodimers**

**A**) A hydrogen bond network forms contacts across the C-terminal half of helices 11 in the dimer interface. From structure of ERα bound to **21**.

**B)** Detail of ligand binding for [F]-AcrEster (**21**) or [F]-Acr (**23**) bound to ERα, showing how the acrylate side chains formed h-bonds with the base of h3 when in the ligand has the F-in conformer. The weaker bond with **21** likely contributes to its higher affinity as it requires less strain on the core (not shown).

**C–D**) Structures of ER with the indicated ligands show multiple ligand binding poses. Compare to **16** in **Fig. 1C** in the F-ring out conformer. **17** showed a mixture of binding modes on one monomer and F-ring out in the other monomer of the dimer.

**E)** Summary of ligand dynamics show how many monomers (out of 4) show the F-ring in or outward facing or have the side chains disordered.

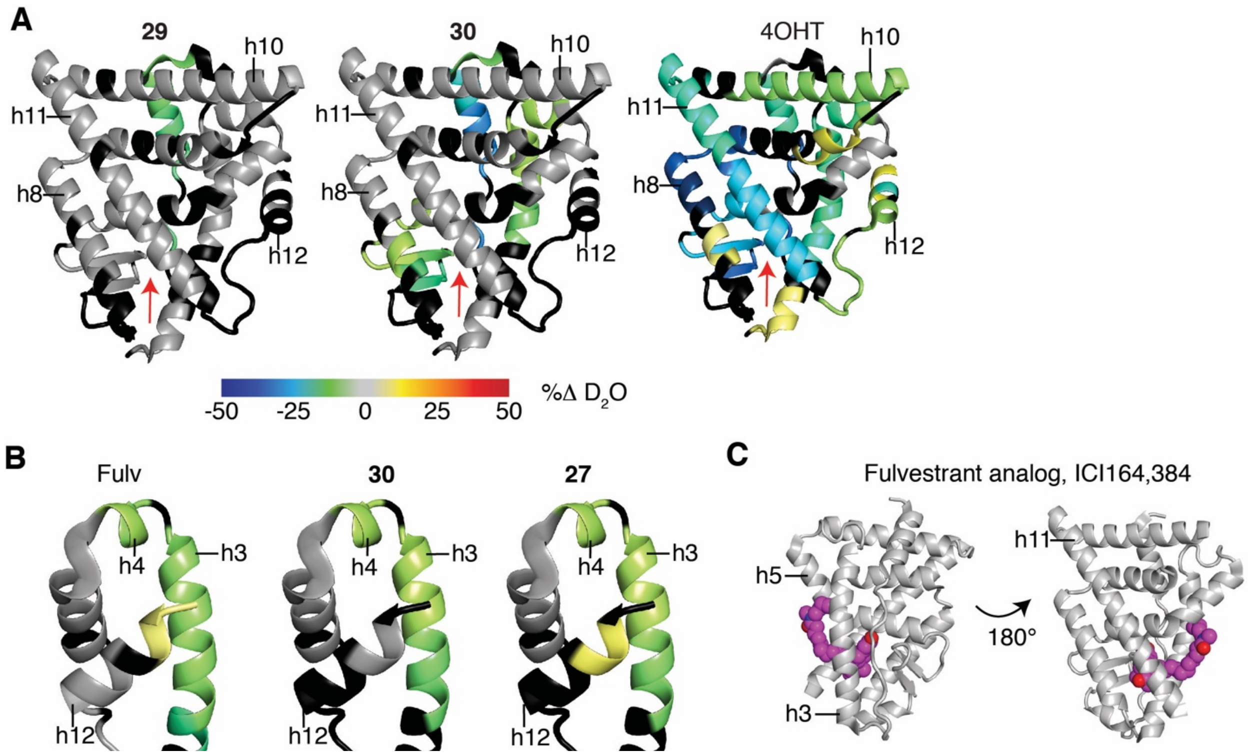


**Figure S4. HDX-MS structural analyses show altered h11/h12 conformations.**

**A–B)** Hydrogen-deuterium exchange mass spectrometry analysis of the Y537S-ERα LBD bound to the indicated ligands. Negative change indicates regions stabilized by ligand compared to apo, and positive change indicates destabilization.

**C)** Structure of ERβ bound to ICI164,384 showed how the side chain docks in the AF-2 surface. From 1HJ1.PDB

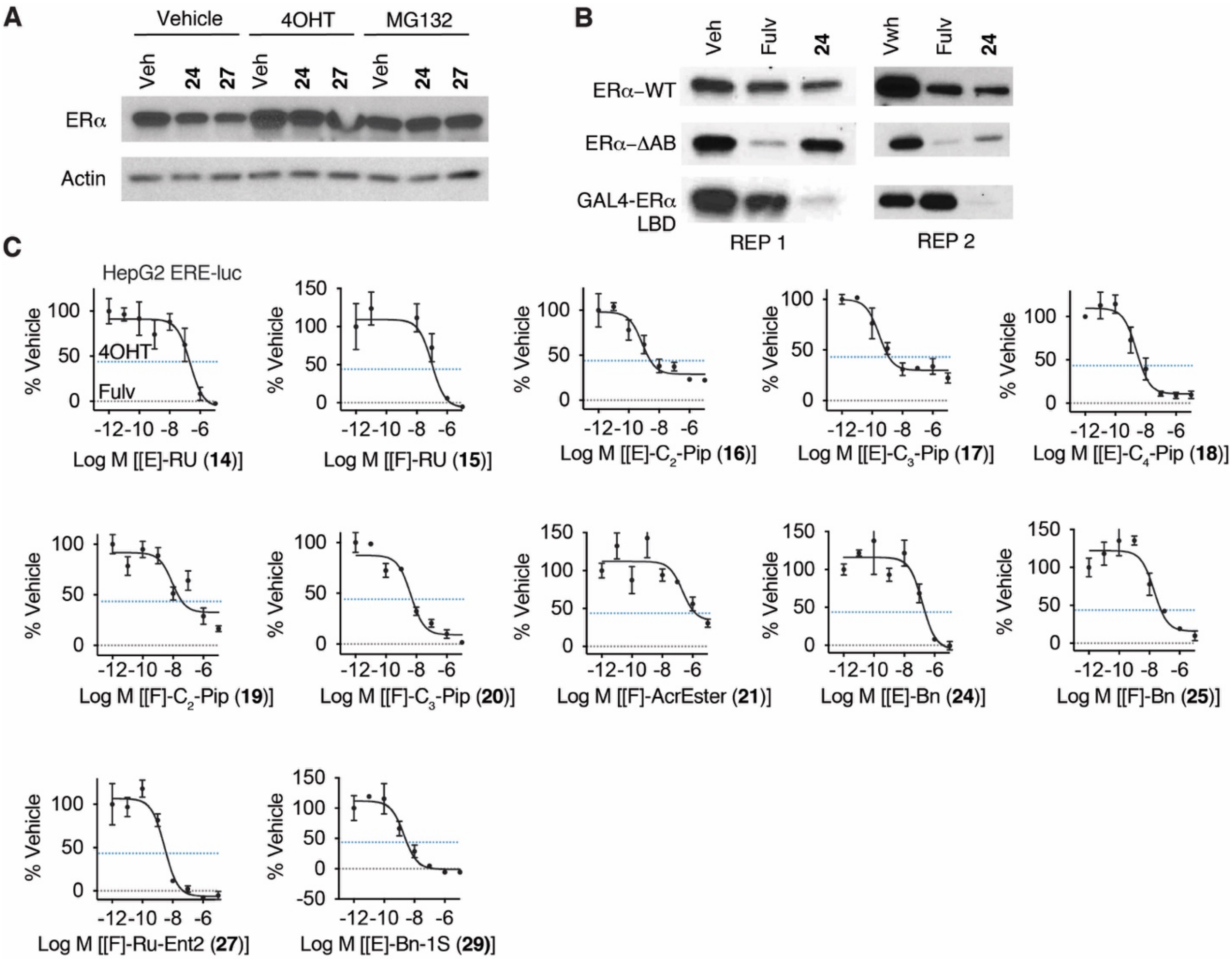


**Figure S5 Characterization of compounds as SERDs or SERMs**

**A**) MCF-7 cells were co-treated with 100 nM of the indicated ligands and 10 nM 4OH-tamoxifen or 30 μM MG132. Whole cell lysates were analyzed by Western blot.

**B**) HepG2 cells were transfected with the indicated ERα expression plasmids and then treated with vehicle, fulvestrant, or 24 for 24 h. Whole cell lysates were analyzed by Western blots. Replicates 1 and 2 are shown.

**C**) HepG2 cells were transfected with ERα expression vector and a 3xERE-luciferase reporter. The next day cells were treated for 24 hrs with the indicated ligands and then processed for luciferase activity. As shown in the first panel, data are normalized to vehicle (100%) and fulvestrant (0%); maximum 4OHT inhibition is indicated by the blue line (ca. 45%). N = 3, mean ± SEM.

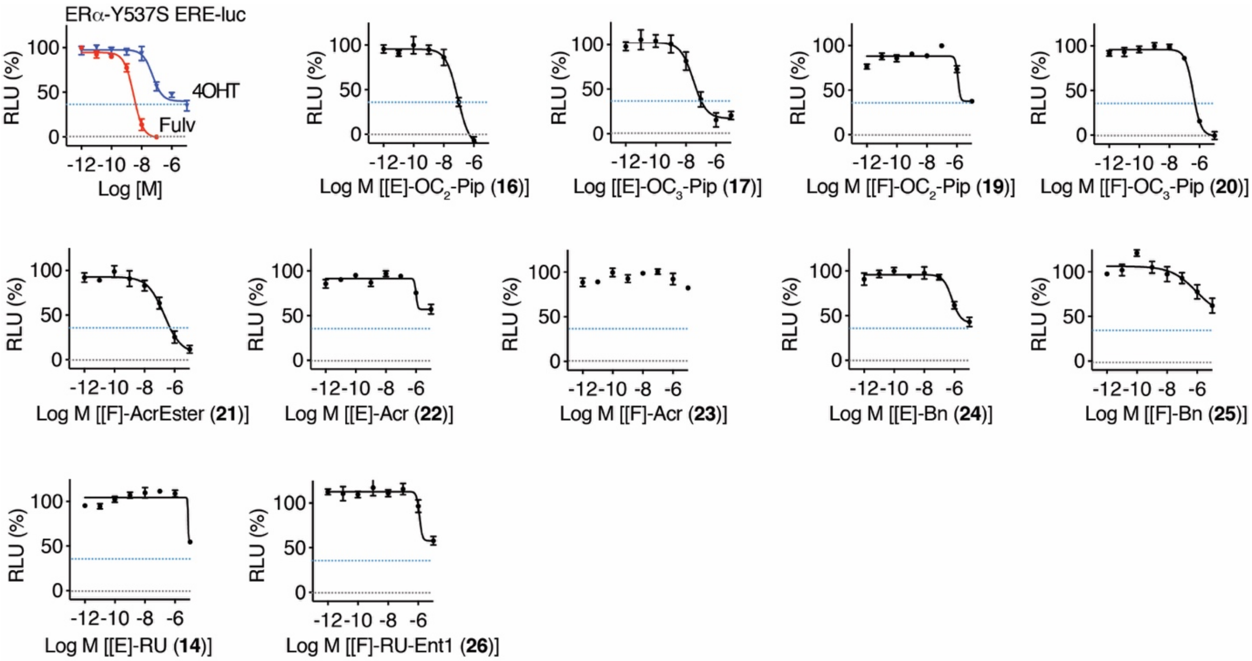


**Figure S6. Activity of DERMIs against the constitutively active mutant ERα-Y537S commonly found in metastatic treatment resistant breast cancer.**

Dose response curves for inhibition of ER-Y537S in 3xERE-luc assay in HEK293T cells. Datapoints are mean ± SEM, N = 3. Blue and black dashed lines indicate the maximal efficacies of 4OHT and Fulv, respectively.

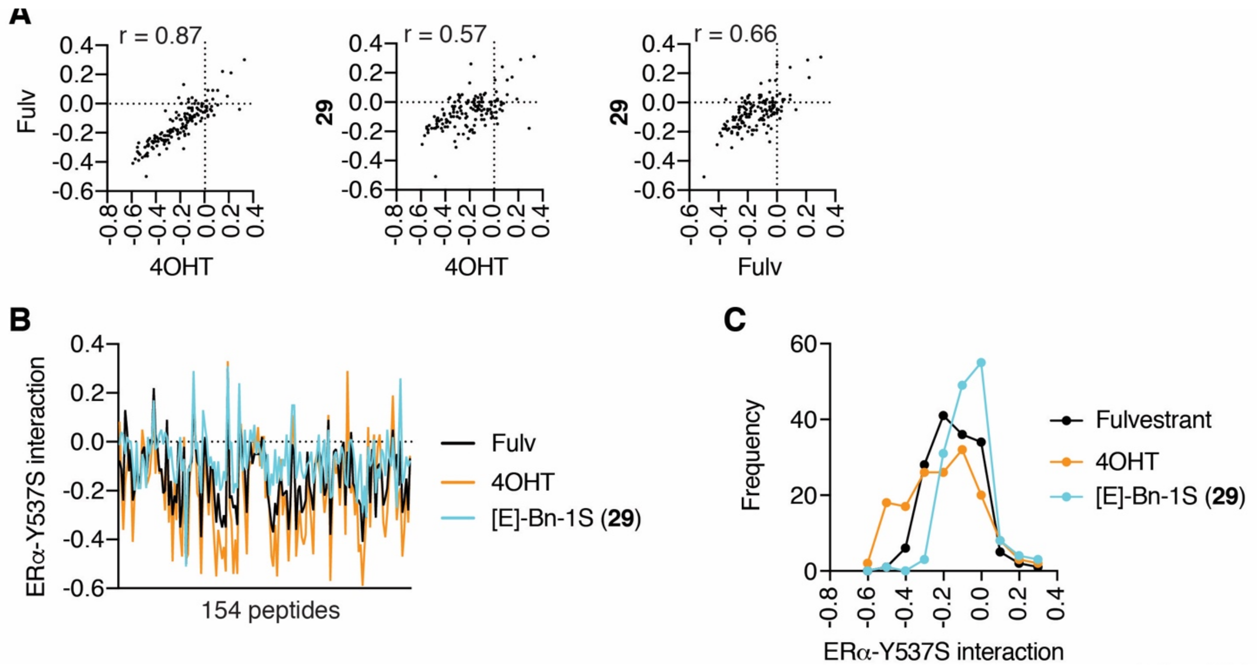


**Figure S7. MARCoNI (Realtime Coregulator-Nuclear receptor Interaction) FRET assay for interaction of full-length ERα-Y537S with 154 peptides derived from nuclear receptor-interacting proteins and the indicated ligands.**

**A)** Pearson correlation for the interaction profiles of the peptides with ERα-Y537S bound to the indicated ligands. Data are log (compound/vehicle).

**B)** Data from the individual ligands is plotted to highlight that **29** showed reduced dismissal of some peptides compared to Fulvestrant and 4OHT.

**C)** Histograph of data from panel **B**.

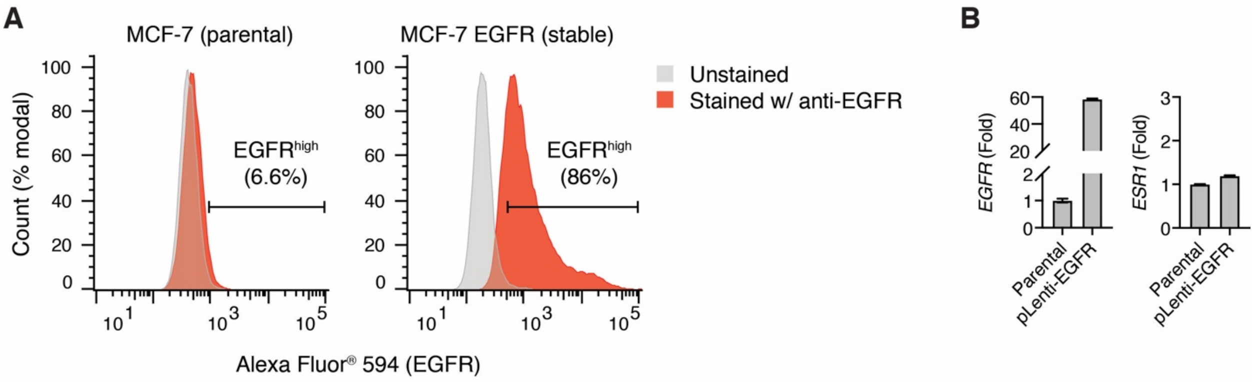


**Figure S8. An EGFR overexpression model of tamoxifen resistance.**

**A)** MCF-7 cells were transduced with a lentivirus expression plasmid for EGFR. Cells were analyzed with FACS for expression of EGFR.

**B)** Parental or EGFR overexpressing MCF-7 cells were analyzed by RT-PCR for expression of the EGFR and ESR1 genes. Data are mean + SEM of two biological replicates

**Supplemental Methods**

**Hydrogen Deuterium Exchange Mass Spectrometry**

Peptide Identification: Peptides were identified using tandem MS (MS/MS) with an Orbitrap mass spectrometer (Q Exactive, ThermoFisher). Product ion spectra were acquired in data-dependent mode with the top five most abundant ions selected for the product ion analysis per scan event. The MS/MS data files were submitted to Mascot (Matrix Science) for peptide identification. Peptides included in the HDX analysis peptide set had a MASCOT score greater than 20 and the MS/MS spectra were verified by manual inspection. The MASCOT search was repeated against a decoy (reverse) sequence and ambiguous identifications were ruled out and not included in the HDX peptide set.

HDX-MS analysis: Protein (10 μM) was incubated with the respective ligands at a 1:10 protein-to-ligand molar ratio for 1 h at room temperature. Next, 5 μl of sample was diluted into 20 μl $D_2O$ buffer (20 mM Tris-HCl, pH 7.4; 150 mM NaCl; 2 mM DTT) and incubated for various time points (0, 10, 60, 300, and 900 s) at 4°C. The deuterium exchange was then slowed by mixing with 25 μl of cold (4°C) 3 M urea and 1% trifluoroacetic acid. Quenched samples were immediately injected into the HDX platform. Upon injection, samples were passed through an immobilized pepsin column (2mm × 2cm) at 200 μl $min^{-1}$ and the digested peptides were captured on a 2mm × 1cm $C_8$ trap column (Agilent) and desalted. Peptides were separated across a 2.1mm × 5cm $C_{18}$ column (1.9 μl Hypersil Gold, ThermoFisher) with a linear gradient of 4% - 40% $CH_3CN$ and 0.3% formic acid, over 5 min. Sample handling, protein digestion and peptide separation were conducted at 4°C. Mass spectrometric data were acquired using an Orbitrap mass spectrometer (Exactive, ThermoFisher). HDX analyses were performed in triplicate, with single preparations of each protein ligand complex. The intensity weighted mean m/z centroid value of each peptide envelope was calculated and subsequently converted into a percentage of deuterium incorporation. This is accomplished determining the observed averages of the undeuterated and fully deuterated spectra and using the conventional formula described elsewhere(1). Statistical significance for the differential HDX data is determined by an unpaired t-test for each time point, a procedure that is integrated into the HDX Workbench software(2). Corrections for back-exchange were made on the basis of an estimated 70% deuterium recovery, and accounting for the known 80% deuterium content of the deuterium exchange buffer.

Data Rendering: The HDX data from all overlapping peptides were consolidated to individual amino acid values using a residue averaging approach. Briefly, for each residue, the deuterium incorporation values and peptide lengths from all overlapping peptides were assembled. A weighting function was applied in which shorter peptides were weighted more heavily and longer peptides were weighted less. Each of the weighted deuterium incorporation values were then averaged to produce a single value for each amino acid. The initial two residues of each peptide, as well as prolines, were omitted from the calculations. This approach is similar to that previously described (3). HDX analyses were performed in triplicate, with single preparations of each purified protein/complex. Statistical significance for the differential HDX data is determined by t test for each time point, and is integrated into the HDX Workbench software (2).

**Luciferase reporter assay**

HepG2 cells were seeded in 10 cm plates containing 10 ml of DMEM (Gibco™ by Thermo Fisher Scientific, cat. no. 11995) supplemented with 10% fetal bovine serum (FBS), 1x GlutaMAX (Gibco™ by Thermo Fisher Scientific, cat. no. 35050061), 1x MEM nonessential amino acids (Corning,cat. no. 25-025-CI), 1x penicillin-streptomycin-neomycin (PSN) antibiotic mixture (Thermo Fisher Scientific, cat. no.15640055), and 2.5 μg/ml Plasmocin™ (Invivogen, cat. no. ant-mp).

The next day, the cells were rinsed with 1x PBS and the medium was replaced with 10 ml of phenol red-free DMEM (Corning, cat. no. 17205CV) supplemented with 10% charcoal-stripped FBS (cs-FBS) (Thermo Fisher Scientific; cat no. A3382101).

The cells were then co-transfected with 5.0 μg of 3xERE-Luc reporter plasmid and 0.5 μg of ERα (WT/mutant) expression plasmid using Fugene HD reagent (Promega, cat no. E2311).

After 24 h, the cells were resuspended in phenol red-free DMEM plus 10% cs-FBS, and transferred to a 384-well plate (Greiner Bio-One, cat. no. 781080) at a density of ~17,000 cells/well containing 25 μl of phenol red-free DMEM plus 10% cs-FBS.

The next day, the test compounds were added using a Biomek NXP 100-nl pintool (Beckman Coulter, Inc.). The plates were sealed with Breathe-Easy permeable membranes (Diversified Biotech, cat. no. BEM-1), covered with stainless steel specimen plate lids (U.S. Patent 6,534,014), and incubated at 37°C overnight. Luciferase activity was measured 24 h later, using the britelite plus reporter gene assay system (PerkinElmer, cat no. 6066761) or the Bright-Glo™ Luciferase Assay System (Promega, cat no. E2620) and an Envision plate reader (PerkinElmer).

**Cell Proliferation Assay**

Cells were suspended in steroid-free media supplemented with 10% charcoal-stripped FBS and passed through a 30-micron strainer (Miltenyl Biotec, cat. no. 130-110-915). 25 µl of the cell suspension (i.e. 1,000 or 2,000 cells) was dispensed into each well of 384-well white, flat-bottom microplates (Greiner Bio-One CellStar, cat no. 781080), using a Martix WellMate Microplate Reagent Dispenser (Thermo Fisher Scientific). The next day, test compounds were added to the wells using a Biomek NXP 100-nl pintool (Beckman Coulter). The plates were sealed with Breathe-Easy permeable membranes (Diversified Biotech, Cat no. BEM-1), covered with a stainless steel specimen plate lid (U.S. Patent 6,534,014), and incubated at 37°C and 5% $CO_2$. “Start plates” with replica wells were stored at -80°C to record the initial number of cells. After 5 days, the start plates were thawed for 15 min at 37°C, and the number of cells/well in all plates were compared. To this end, 25 µl of CellTiter-Glo® assay reagent (Promega, cat no. G7573) was added to each well using the Wellmate reagent dispenser. The plates were shaken gently at room temperature for 5 min on an orbital shaker, and then allowed to sit for 5 min. Luminescence was measured using an Envision plate reader (PerkinElmer). Proliferation data was normalized using the initial number of cells as 0%, and the final number of cells in vehicle (DMSO)-treated wells as 100%.
